# Genome assembly and annotation of the mermithid nematode *Mermis nigrescens*

**DOI:** 10.1101/2022.11.05.515230

**Authors:** Upendra R. Bhattarai, Robert Poulin, Neil J. Gemmell, Eddy Dowle

## Abstract

Genetic studies of nematodes have been dominated by *Caenorhabditis elegans* as a model species. Lack of genomic resources has been a limiting factor for expansion of genetic research to other groups of nematodes. Here, we report a draft genome assembly of a mermithid nematode, *Mermis nigrescens*. Mermithidae are insect parasitic nematodes with hosts including a wide range of terrestrial arthropods. We sequenced, assembled, and annotated the whole genome of *M. nigrescens* using nanopore long-reads and 10X chromium link-reads. The assembly is 524 Mb in size consisting of 867 scaffolds. The N50 value is 2.42 Mb, and half of the assembly is in the 30 longest scaffolds. The assembly BUSCO score from the eukaryotic database (eukaryota_odb10) indicates that the genome is 86.7% complete and 5.1% partial. The genome has a high level of heterozygosity (6.6%) with a repeat content of 78.7%. mRNA-seq reads from different sized nematodes (≤2 cm, 3.5-7 cm, and >7 cm body length) representing different developmental stages were also generated and used for the genome annotation. Using ab initio and evidence-based gene model predictions, 12,313 protein-coding genes and 24,186 mRNAs were annotated. These genomic resources will help researchers investigate the various aspects of the biology and host-parasite interactions of mermithid nematodes.

## Introduction

Nematodes are a highly diverse group with over one million estimated species (Scheffers et al. 2012; Smythe et al. 2019). They are ubiquitous, occurring in habitats ranging from deep oceans to mountain peaks (Schratzberger et al. 2019). Most animals and plants host at least one species of parasitic nematode (Blaxter and Koutsovoulos 2015). Despite representing an important component of all natural ecosystems (Ferris 2010; Cardoso et al. 2016), most nematode species remain undescribed (Dobson et al. 2008).

Nematodes fall into two classes based on both molecular and morphological systematics: Chromadorea and Enoplea (Blaxter et al. 1998; Liu et al. 2013). They have a conserved body plan with similar structural organisation and cellular morphologies (Basyoni and Rizk 2016). However, comparative analyses have highlighted the molecular and physiological diversity within the phylum (Mitreva et al. 2005; Coghlan et al. 2019). *Caenorhabditis elegans* is a Chromadorean that has been at the forefront of genetic research as a model species. In contrast, there exists very little genomic information from the members of class Enoplea. Most studies on Enoplea focus on *Trichinella spiralis* (Enoplea: Trichocephalida) because of its importance as a mammalian parasite (Mitreva et al. 2011). There are also several insect parasitic taxa within the Enoplea, but only the genome of *Romanomermis culicivorax* (Dorylaimia: Mermithidae) is publicly available (Schiffer et al. 2013). The genomes of *T. spiralis* and *R. culicivorax* differ significantly in size (64Mb vs 320Mb) and other attributes like the proportion of repetitive sequences and transposable elements, highlighting the genetic diversity within the Enoplea (Schiffer et al. 2013).

Mermithids are obligate endoparasitic nematodes. They represent a large portion of the understudied nematode class Enoplea. There are over 100 described mermithid species; however, their biology and distribution are poorly studied (Poinar 1975; Presswell et al. 2015). Past studies have paid special attention to mermithid species that have potential as biological control agents against insect pests (Petersen 1973; Abagli et al. 2019). They have adapted to a wide range of arthropod hosts, from terrestrial to aquatic taxa (Yeates and Buckley 2009; Zamani 2014; Kubo et al. 2016; Tong et al. 2021). Of the few well-studied species, many have broad geographical distributions. For example, *Isomermis* species infect several Simuliid hosts throughout Africa, Europe, and North America (Gradinarov 2014). Similarly, *Mermis nigrescens* has been reported from different parts of Europe, Asia, North and South America, Australia, and New Zealand, infecting various hosts across its broad range (Presswell et al. 2015).

Furthermore, several mermithids manipulate their host’s behaviour for their own benefit (Herbison et al. 2019). Adult mermithids are free-living; after completion of their development within an arthropod host, they must emerge from the host to mate and lay eggs. As they are prone to desiccation, they must emerge in water or water-saturated substrate. Since many mermithids parasitize terrestrial hosts, the need to emerge in water has selected for mermithids capable of inducing water-seeking behaviour in their host. For example, the mermithid nematode *Strelkovimermis spiculatus* modifies the behaviour of its adult female mosquito host to make it seek water instead of a blood meal, providing a dispersal means for the nematodes and ensuring their return to a suitable habitat for reproduction (Allahverdipour et al. 2019). Similarly, *M. nigrescens* induces water-seeking behaviour in its terrestrial earwig host (*Forficula auricularia*), enabling the nematodes to emerge into a favourable environment to continue their life cycle (Presswell et al. 2015; Herbison et al. 2019).

Here, we present a high-quality genome assembly of *Mermis nigrescens*. The life cycle, morphological, and molecular characteristics of *M. nigrescens* have been well studied (Baker and Capinera 1997; Presswell et al. 2015). However, the lack of whole-genome information greatly limits the genetic investigations of its biology and adaptations, such as its ability to alter host behaviour. This genome will also fill the knowledge and resource gap for Enoplean genomes. It will further facilitate the application of a range of molecular tools and approaches to study the genetic underpinnings of the wide geographical distribution and successful host exploitation of mermithid nematodes.

## Methods

### Sampling and Nucleotide extractions

Nematodes (*Mermis nigrescens*) were dissected out of the European Earwigs (*Forficula auricularia*) collected from the Dunedin Botanic Garden (Latitude: -45° 51’ 27.59” S; Longitude: 170° 31’ 15.56” E) and reared in a temperature-controlled room (Temperature: cycling from 15 to 12 °C, day/night; Photoperiod of L:D 16:8) in the Department of Zoology, University of Otago, Dunedin. Nematodes thus obtained were snap-frozen in liquid nitrogen and stored at -80°C until further use. Individual nematodes were used for each DNA extraction. Individuals were not sexed prior to sequencing due to the difficulty in achieving this in a manner timely enough to maintain high-quality RNA and DNA samples. DNA extracted using DNeasy® Blood & Tissue Kit (Qiagen, USA) was used for nanopore sequencing. DNA extracted using Nanobind Tissue Big DNA kit (Circulomics, USA) was used for 10x sequencing library preparation. RNase treatment to remove RNA was performed using 4μl of RNase A (10mg/ml) per 200μl of template following DNA extraction. DNA samples were quantified and quality assessed using a Qubit 2.0 Fluorometer (Life Technologies, USA) and Nanodrop. High-quality DNA samples were stored at -20°C and were used within a week of extraction.

Total RNA from different sized nematodes (≤2cm, 3.5-7cm, and >7cm body length) representing different developmental stages was extracted using Direct-zol™ RNA MicroPrep kit (Zymo Research) following the manufacturer’s instruction and stored at -80°C until further use. RNA from five individuals from each size category (small, medium, and large) were individually extracted and later pooled at equimolarity to make a single sequencing library for each category. Samples were quantified and quality checked on a Qubit 2.0 Fluorometer (Life Technologies, USA) and Nanodrop. RNA integrity for the pooled samples was assessed using a Fragment Analyzer (Advanced Analytical Technologies Inc., USA) at the Otago Genomics Facility (OGF), University of Otago, Dunedin, New Zealand. The RNA quality number (RQN) values ranged from 8.3 to 9.5, indicating high-quality samples.

### Library preparation and sequencing

A long read sequencing library for Oxford minion was prepared with 416.5 ng of genomic DNA using a ligation sequencing kit (SQK-LSK109) (Oxford Nanopore Technologies, Oxford, UK) following the manufacturer’s instructions. Lambda phage (DNA CS) was used as a positive control. The prepared library was sequenced with R9 chemistry MinION flow cell (FLO-MIN106) (Oxford Nanopore Technologies, UK).

A linked read library was prepared at the Genetic Analysis Services (GAS), University of Otago (Dunedin, New Zealand). DNA was size-selected for fragments over 40 Kbp using a Blue Pippin (Sage Science, USA). A chromium 10x linked read (10x Genomics, USA) library was prepared following the manufacturer’s instructions. The library was sequenced on the Illumina Nova-seq platform to generate paired-end reads (2×151bp) at the Garvan Institute, Australia.

TruSeq stranded mRNA libraries were prepared and sequenced in OGF as 2×100 bp paired-end reads across two Rapid V2 flowcell lanes of HiSeq 2500.

### Genome size estimation

Genome size was estimated with flow cytometry and a genomic k-mer based approach. Flow cytometry was carried on two whole worms at Flowjoanna (Palmerston North, NZ, USA). The sample preparation protocol for the flowcytometry is described in our earlier study (Bhattarai et al. 2022). Rooster red blood cells (RRBC) from the domestic chicken (*Gallus gallus*) was used a reference. Samples were then processed on a FACSCalibur (BD Biosciences, USA) and the data were analyzed using Flowjo (BD Biosciences, USA).

The short-read data from the 10x Chromium library were used for the k-mer based approach. We used scaff_reads script from Scaff10x (v.5.0) to remove the 10x link adapters and low-quality reads, while any remaining Illumina adapters were removed using Trimmomatic (v.0.39) with options: LEADING:3 TRAILING:3 SLIDINGWINDOW:4:15 MINLEN:35. A k-mer size of 21 was used with KMC (v.3.1.1) (Kokot et al. 2017) to produce a histogram that was visualised in the Genomescope (v.2.0) web browser.

### Bioinformatic pipeline

We used various software and packages to assemble and refine the genome (Figure 1). The pipeline and the scripts used in this project are available on GitHub (https://github.com/upendrabhattarai/Nematode_Genome_Project). The bioinformatics software and packages were run in New Zealand eScience Infrastructure (NeSI).

**Figure 1:**
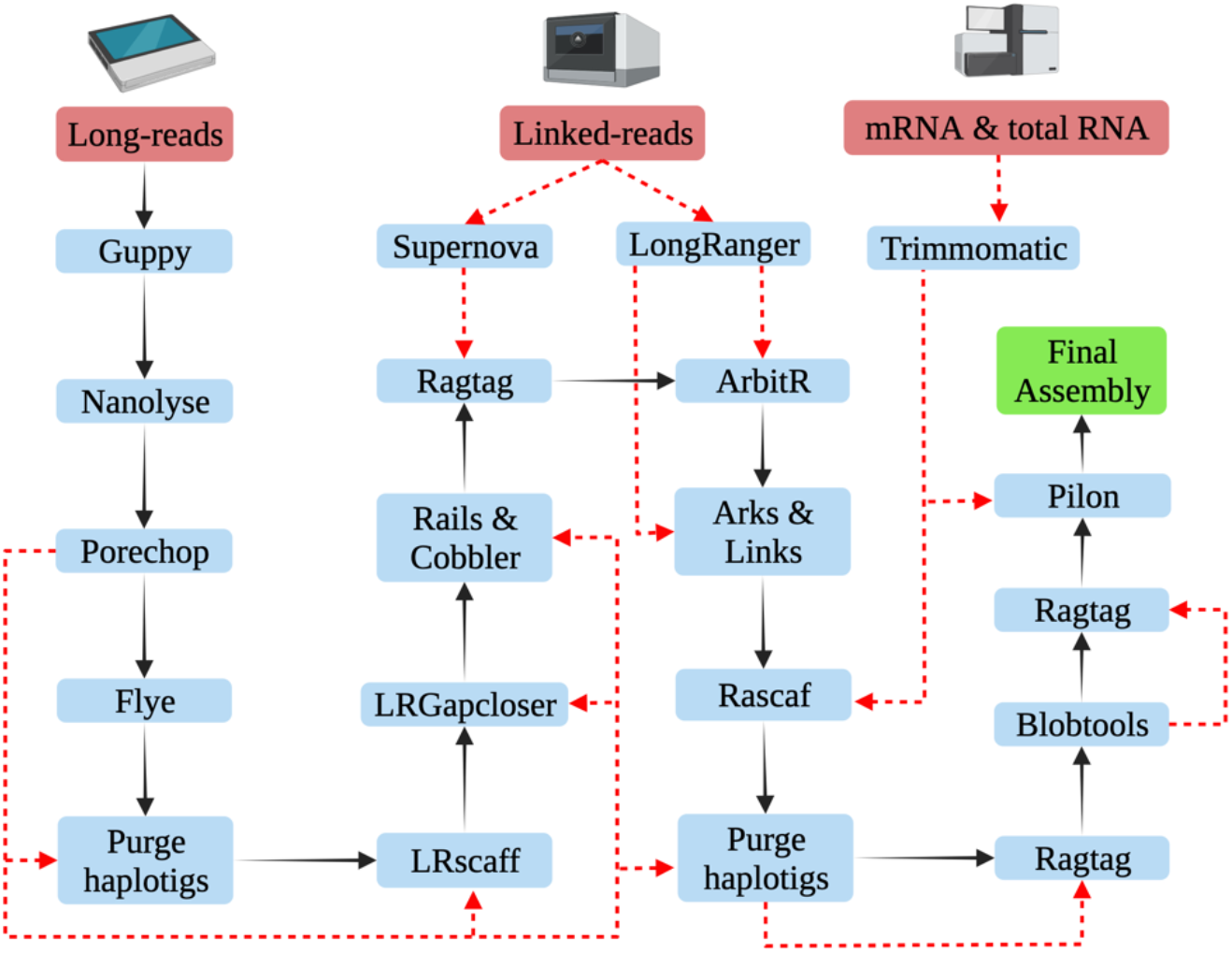
Schematic representation of the assembly pipeline for the *Mermis nigrescens* genome. The black arrows represent the workflow and the red dotted lines represent the input data in the pipeline (Created with BioRender.com).

### Genome and transcriptome assembly

All the Nanopore reads were base called using Guppy (v.5.0.7) and assembled using Flye (v.2.9) (Kolmogorov et al. 2019) with default parameters. The paired end reads from the chromium library were assembled using supernova (Weisenfeld et al. 2017). BUSCO (v.4.1.4) (Simão et al. 2015) and Quast (v.5.0.2) (Gurevich et al. 2013) were used to evaluate the assembly statistics. The assembly from long reads outperformed the linked reads for the BUSCO completeness and assembly contiguity. Therefore, long read assembly was used as a primary assembly, and the linked read assembly was used for scaffolding later in the pipeline (Figure 1). Multiple scaffolding and gap-closing steps were performed using Lrscaf (v.1.1.11) (Qin et al. 2019), Rails (v.1.5.1) and Cobbler (v.0.6.1) (Warren 2016), Ragtag (v.2.1.0) (Alonge et al. 2019), ArbitR (v.0.2) (Hiltunen et al. 2021), Arks (v.1.0.4) (Coombe et al. 2018), Links (v.1.8.7) (Warren et al. 2015) and Lrgapcloser (Xu et al. 2019). mRNA-seq reads obtained for genome annotation purposes, and total RNA-seq reads sequenced for another project (manuscript under preparation) were also used for scaffolding the assembly with Rascaf (Song et al. 2016). Purgehaplotigs (Roach et al. 2018) removed haplotigs from the assembly. Removed haplotig sequences were used to further scaffold the assembly with Ragtag (v.2.1.0). Blobtools2 (Laetsch and Blaxter 2017) was used to remove and discard bacterial reads and remove low coverage (<5x coverage) and short (<1000bp) scaffolds. Low coverage and short scaffolds were then used to scaffold the filtered assembly with Ragtag (v.2.1.0) (Alonge et al. 2019). Finally, Pilon (v.1.24) was used to polish the assembly using mRNA-seq data.

The adapters and low quality reads from mRNA-seq data were filtered using Trimmomatic (v.0.38) (Bolger et al. 2014) with options HEADCROP:15 LEADING:3 TRAILING:3 SLIDINGWINDOW:4:20 MINLEN:35. Clean data was de novo assembled with Trinity (v.2.13.2) (Grabherr et al. 2011) using all the default parameters.

### Repetitive content masking

A custom repeat library was generated using de novo and homology-based identifiers, including LTRharvest (Ellinghaus et al. 2008), LTRdigest (Steinbiss et al. 2009), RepeatModeler (Flynn et al. 2020), TransposonPSI (Brian Haas 2010), SINEBase (Vassetzky and Kramerov 2013), and MITE-Tracker (Crescente et al. 2018). Individual libraries from each software were concatenated, and sequences with more than 80% similarity were merged to remove redundancy using usearch (v.11.0.667) (Edgar 2010). The library was then classified using RepeatClassifier. Sequences with unknown categories in the library were mapped against the reviewed nematode database from UniProtKB/Swiss-Prot database (e-value <1e-01); if not annotated as repeat sequences, they were removed from the library. The final repeat library was used in RepeatMasker (v.4.1.0) (Chen 2004) to mask the repeats and generate a report. The repeat library was used as an input to the MAKER2 pipeline (Holt and Yandell 2011).

### Genome annotation

The assembly was annotated using evidence-based and ab initio gene model predictions through MAKER2 (v.2.31.9) (Holt and Yandell 2011) and Braker (v.2.16) (Hoff et al. 2019) pipelines. The first round of MAKER2 was run with 180,630 mRNA transcripts *denovo* assembled with the Trinity pipeline (Grabherr et al. 2011) along with 511,117 mRNA and 1,148,233 protein sequences from all the Dorylaimia species available in NCBI and WormBase (https://wormbase.org) databases. mRNA-seq reads and output from GenMark-ES were used to train Augustus with the Braker pipeline. SNAP was trained after each round of MAKER2. Trained SNAP and Augustus were used for two more iterations of the MAKER2 pipeline.

We used InterProScan (v.5.51-85.0) (Jones et al. 2014) for the functional annotation on the predicted protein sequences from MAKER and retrieved InterPro ID, PFAM domains, and Gene Ontology (GO) terms. Furthermore, the Uniprot database with BLAST was used to assign gene descriptors to each transcript based on the best hit.

## Results and Discussion

### Genome size estimates

The flowcytometry estimate of the nematode genome size is 814.185±24.2 Mb (mean ±SD), however, the nematode samples yielded very low nuclei counts, i.e. as low as <200 total nuclei for one of the samples. Given that this is a whole animal digest and the very low nuclei counts in the samples, the calculated pg/nucleus may not be accurate. To get better genome size estimates by flowcytometry in this nematode, we recommend increasing the sample size and replicated measurements from the nuclei suspension in future experiments.

The k-mer based analysis estimated the genome size to be 336.07 Mb with 6.56% of heterozygosity (Figure 2). A high level of heterozygosity can result in genome assemblers generating multiple copies of the region, leading to larger genome size and reduced contiguity (Guan et al. 2020; Asalone et al. 2020). This may explain why the assembled genome size (524.2 Mb) is substantially larger than the k-mer estimates. Furthermore, as *M. nigrescens* can also reproduce parthenogenetically(Christie J. R. 1937), we assume that the high level of heterozygosity in their genome is associated with their parthenogenetic mode of reproduction. Similarly, a highly heterozygous genome (5.7%) was also reported for another parthenogenetic nematode, *Diploscapter coronatus* (Hiraki et al. 2017).

**Figure 2:**
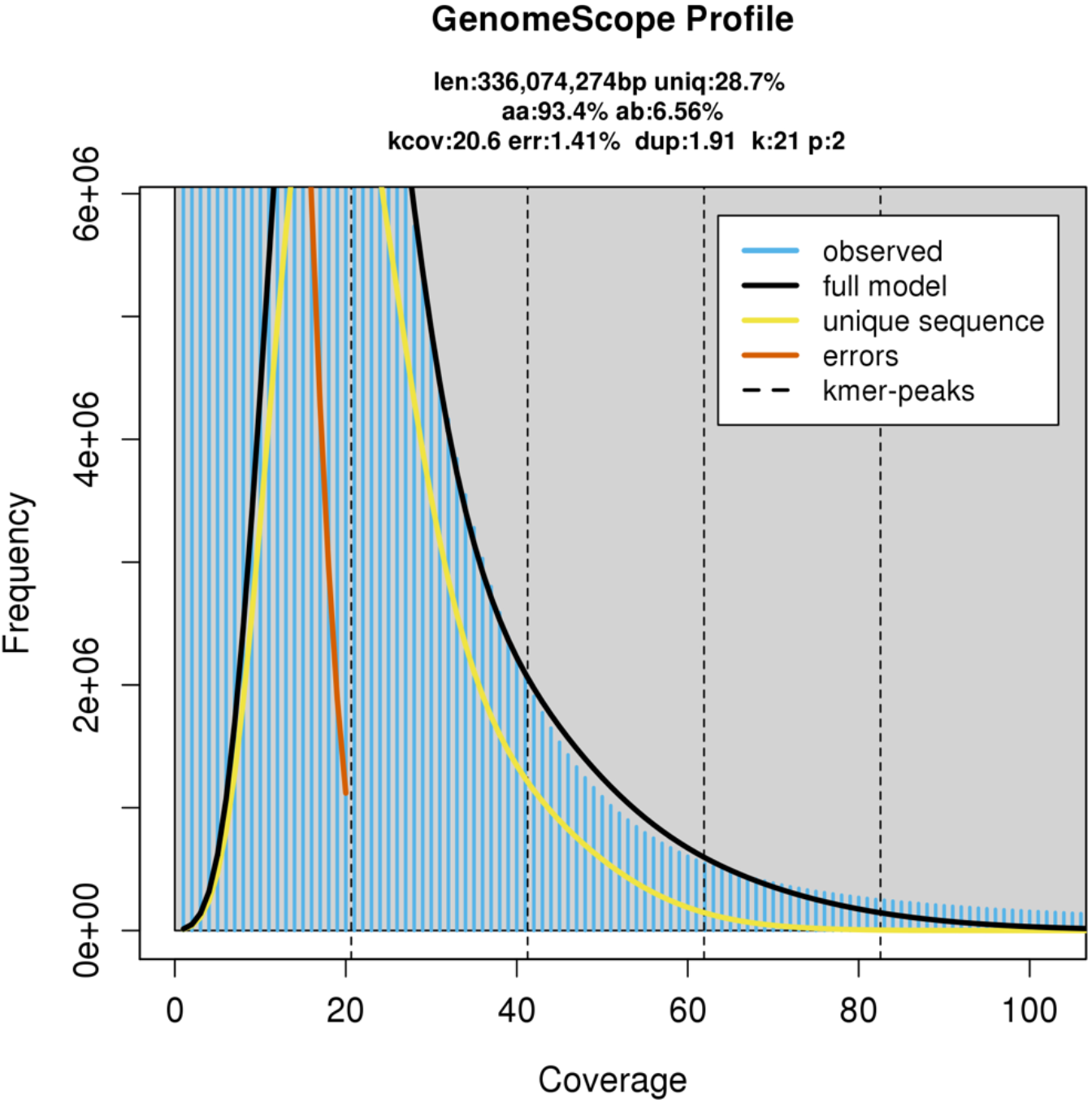
The k-mer analysis of *Mermis nigrescens* genome using short reads. A k-mer size of 21 was used in KMC (v.3.1.1) to produce a histogram and was visualised in the Genomescope (v.2.0) web browser. It shows a genome size of 336.07 Mb with heterozygosity of 6.56%.

### Genome and transcriptome assembly

A total of 6.9 Gb of sequencing data was generated from a Nanopore minion flowcell run consisting of 530,888 reads with an N50 length of 17,849 bp (Table 1). Flye (v.2.7.1) produced the primary long read-assembly with 16,414 contigs with N50 of 94,974 bp and a total length of ∼714 Mb. Quast reported 88.45% complete BUSCO from the Eukaryote database for this assembly.

**Table 1.** Sequencing output from Oxford nanopore minion.

| Reads | Bases | Median Read Length | N50 length | Median Read Quality |
| --- | --- | --- | --- | --- |
| 530,888 | 6,935,586,657 | 11,833 | 17,849 | 13.63 |

The 10x chromium library yielded 450.3 million paired-end reads. These linked reads with the Supernova assembler produced an assembly of 582.5 Mb with 147,205 scaffolds and N50 of 9,664 bp. Quast reported only 20.13% of complete BUSCO genes for this assembly (Table 2). Comparing the contiguity and completeness of these two assemblies, we used Flye assembly as a primary assembly and the supernova assembly was used in scaffolding steps later in the pipeline (Figure 1).

**Table 2.**
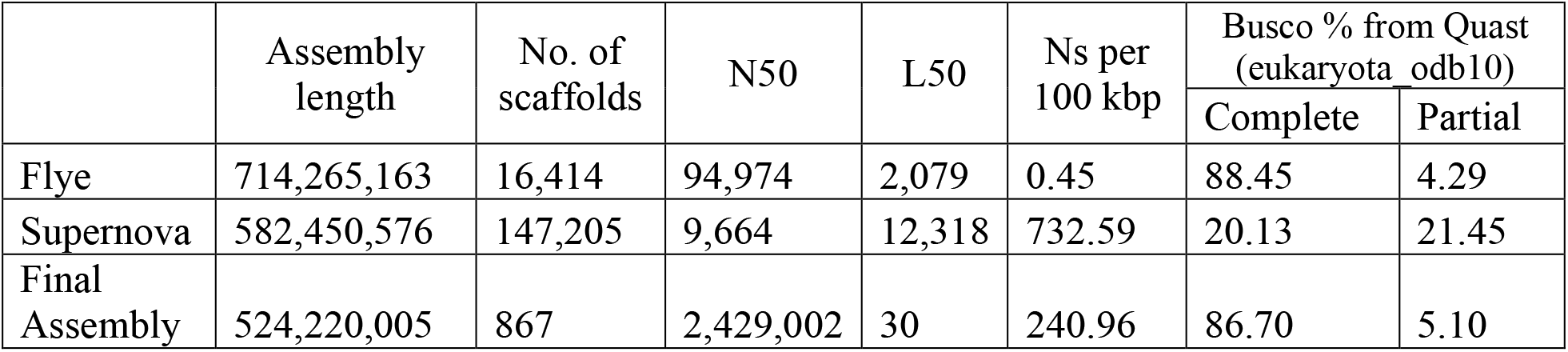
Assembly statistics for the genome assembly of *Mermis nigrescens*.

The assembly statistics after each round of processing are presented in Table 3. The final assembly has a size of 524.2 Mb with 867 scaffolds. The assembly has the N50 of 2,429,002 bp, and 50% of the assembly is in the 30 longest scaffolds. The k-mer based analysis estimated the genome size to be 336 Mb with a heterozygosity of 6.56%.. Gaps in the assembly account for 240.97 bp per 100 Kbp length. The final assembly has a complete BUSCO score of 86.7% and a partial BUSCO score of 5.1% to the eukaryotic database and (Figure 3A).

**Table 3.** Assembly statistics after each round of processing.

| Assembly steps | Assembly length | No. of scaffolds | N50 | L50 | Ns per 100 kbp | Busco % from Quast (eukaryota odb10) |  |
| --- | --- | --- | --- | --- | --- | --- | --- |
|  |  |  |  |  |  | Complete | Partial |
| Flye | 714,265,163 | 16,414 | 94,974 | 2,079 | 0.45 | 88.45 | 4.29 |
| Purgehaplotigs | 547,497,121 | 8,101 | 125,056 | 1,325 | 0.57 | 85.15 | 7.59 |
| Lrscaff | 739,840,003 | 5,500 | 263,548 | 890 | 2122.06 | 88.45 | 3.96 |
| LRGapcloser | 739,845,718 | 5,500 | 263,608 | 890 | 283.38 | 88.45 | 3.96 |
| Rails & Cobbler | 740,070,390 | 5,454 | 266,049 | 885 | 256.75 | 89.11 | 3.30 |
| Ragtag | 740,076,890 | 5,389 | 266,578 | 883 | 257.63 | 89.11 | 3.30 |
| ArbitR | 738,644,384 | 4,958 | 308,460 | 738 | 262.84 | 89.11 | 3.30 |
| ARKS & LINKS | 738,665,484 | 4,747 | 328,769 | 687 | 265.69 | 88.45 | 3.96 |
| Rascaf | 738,576,542 | 4,343 | 388,132 | 575 | 253.68 | 88.45 | 3.96 |
| Purgehaplotigs | 587,968,296 | 3,113 | 443,562 | 412 | 244.20 | 86.14 | 5.61 |
| Ragtag | 588,012,496 | 2,671 | 533,508 | 328 | 251.70 | 86.14 | 5.61 |
| Blobtools | 524,163,746 | 1541 | 553,974 | 285 | 228.13 | 80.86 | 5.94 |
| Ragtag | 524,231,146 | 867 | 2,428,984 | 30 | 240.95 | 80.86 | 6.27 |
| Pilon | 524,220,005 | 867 | 2,429,002 | 30 | 240.96 | 86.70 | 5.10 |

**Figure 3:**
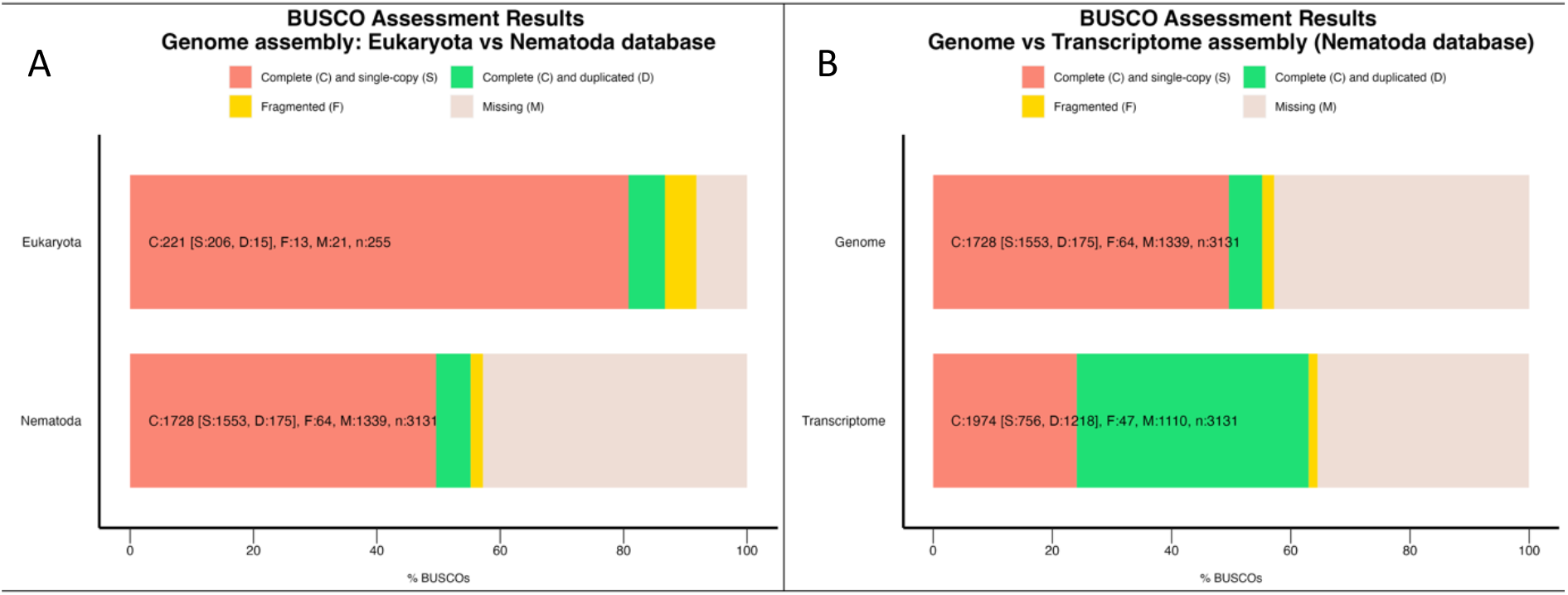
The BUSCO (v.4.1.4) analysis report for the genome and transcriptome assembly of *Mermis nigrescens*. A) Comparison of the BUSCO scores from the Eukaryote (eukaryota_odb10) and the Nematode (nematoda_odb10) database for the genome. B) Comparison of BUSCO score for the genome and transcriptome assemblies with the nematoda_odb10 database.

mRNA-seq libraries generated 288.9 million paired-end reads. They were *de novo* assembled using Trinity (v.2.13.2) (Grabherr et al. 2011) producing 180,630 transcripts with N50 of 1,362 bp including 103,144 trinity genes. The Busco score from the nematoda_odb10 database was 63% for the transcriptome assembly and 55.2% complete for the genome assembly (Figure 3B). This suggests that the nematode Busco database (nematoda_odb10) does not provide a good representation of the *M. nigrescens* genome, further highlighting the high genetic diversity among nematodes.

*Romanomermis culicivorax* is the only mermithid nematode with a publicly available whole-genome assembly (Schiffer et al. 2013) to date. Its genome assembly has a 322 Mb size with N50 of 17,632 bp. It has 62,537 scaffolds and 37.3% complete Busco genes (nematoda_odb10, n=3131). Between these two assemblies, the *M. nigrescens*’ genome assembly is more contiguous and complete and can be used as a better representative of mermithid nematodes.

### Genes and repeats annotation

A total of 12,313 protein coding genes and 24,186 mRNAs were identified in the genome. The mean gene length is 12,224 bp, and the total gene length is 150.5 Mb. The longest gene annotated is 206,129 bp (Table 4). Functional annotation resulted in 7,496 genes annotated to the InterPro and Pfam databases and 4,623 annotated to Gene Ontology terms (Table 5). Quality of annotation was measured using Annotation Edit Distance (AED) score, and 96% of the annotated genes have an AED score of 0.5 or less, ensuring highly confident gene prediction (Figure 4).

**Table 4.** Genome annotation summary for *Mermis nigrescens*.

|  |  |
| --- | --- |
| Number of genes | 12,313 |
| Number of mRNAs | 24,186 |
| Number of exons | 152,566 |
| Number of introns | 1,073,659 |
| Number of CDS | 24,185 |
| Total gene length | 150,517,544 |
| Total mRNA length | 298,339,285 |
| Total exon length | 30,891,521 |
| Total intron length | 2,450,417,037 |
| Total CDS length | 23,760,135 |
| Longest gene | 206,129 |
| Longest mRNA | 206,129 |
| Longest exon | 6,519 |
| Longest intron | 539,449 |
| Longest CDS | 19,164 |
| mean gene length | 12,224 |
| mean mRNA length | 12,335 |
| mean exon length | 202 |
| mean intron length | 2,282 |
| mean CDS length | 982 |

**Table 5.** Functional annotation statistics from different databases.

| Database | No. of terms linked to mRNA | No. of mRNA with term | No. of gene with term |
| --- | --- | --- | --- |
| InterPro | 21,702 | 16,007 | 7,496 |
| Gene Ontology | 20,530 | 10,180 | 4,623 |
| Pfam | 21,954 | 16,007 | 7,496 |

**Figure 4:**
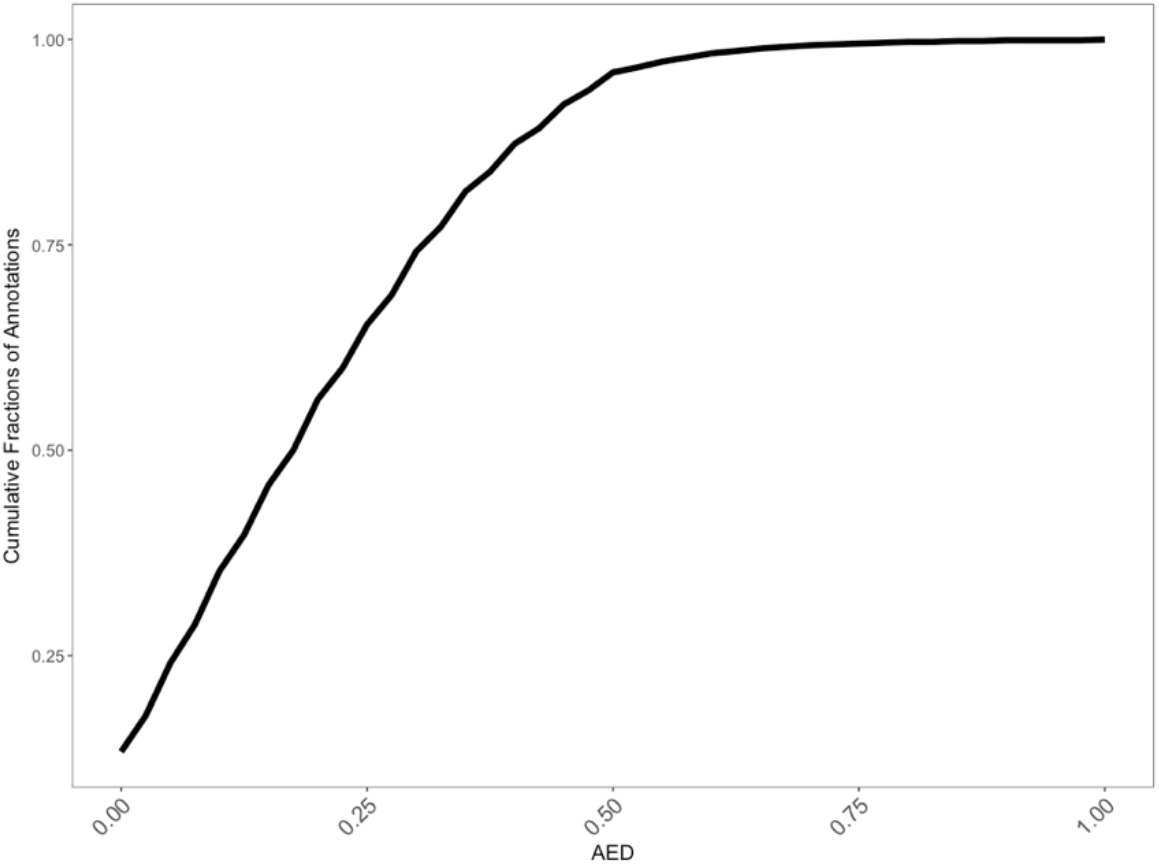
Annotation quality with AED scores. The y-axis shows the cumulative fractions of annotations and the x-axis their AED scores.

The repeat contents in the genome amount to 412.78 Mb comprising 78.74% of the whole assembly. It includes 17.86% of retroelements, 21.97% of DNA transposons, 1.09% rolling circles and 37.18% unclassified repeatomes (Table 6). Compared to other nematode species, this is a very high proportion of repeats in the genome. For example, the repeat contents in *R. culcivorax* genome is 47% (Schiffer et al. 2013), in *Trichinella spiralis* genome it is 19.8% (Mitreva et al. 2011) and in *Caenorhabditis elegans* genome it is 17% (*C elegans* Sequencing Consortium 1998). At present, we do not know the significance of such a highly repetitive genome in *M. nigrescens*. However, further investigations might shed light on its biological and evolutionary significance.

**Table 6.** Repeat content analysis in *Mermis nigresens* genome assembly.

|  |  |  |  |  |
| --- | --- | --- | --- | --- |
| No. of sequences: | 867 |  |  |  |
| Total length (bp): | 524,220,005 |  |  |  |
| GC level: | 36.63% |  |  |  |
| Bases masked: | 412,783,310 bp (78.74%) |  |  |  |
|  |  | Numbers* | Length (bp) | Percentage |
| Retroelements |  | 305,036 | 93,643,203 | 17.86% |
|  | SINEs: | 2,942 | 369,353 | 0.07% |
|  | Penelope | 25,117 | 7,072,788 | 1.35% |
|  | LINEs: | 147,865 | 40,170,331 | 7.66% |
|  | CRE/SLACS | - | - | 0.00% |
|  | L2/CR1/Rex | 85,577 | 23,760,597 | 4.53% |
|  | R1/LOA/Jockey | 80 | 54,133 | 0.01% |
|  | R2/R4/NeSL | 166 | 29,796 | 0.01% |
|  | RTE/Bov-B | 13,943 | 3,447,682 | 0.66% |
|  | L1/CIN4 | 13,163 | 2,091,210 | 0.40% |
|  | LTR elements: | 154,229 | 53,103,519 | 10.13% |
|  | BEL/Pao | 8,192 | 2,959,770 | 0.56% |
|  | Ty1/Copia | 731 | 84,377 | 0.02% |
|  | Gypsy/DIRS1 | 98,159 | 41,038,229 | 7.83% |
|  | Retroviral | 17,348 | 3,301,532 | 0.63% |
| DNA transposons |  | 503,758 | 115,186,810 | 21.97% |
|  | hobo-Activator | 105,906 | 20,614,222 | 3.93% |
|  | Tc1-IS630-Pogo | 36,016 | 7,078,658 | 1.35% |
|  | En-Spm | - | - | 0.00% |
|  | MuDR-IS905 | - | - | 0.00% |
|  | PiggyBac | 1,737 | 704,735 | 0.13% |
|  | Tourist/Harbinger | 25,116 | 5,166,803 | 0.99% |
|  | Other (Mirage,<br>P-element,<br>Transib) | - | - | 0.00% |
| Rolling-circles |  | 28,792 | 5,739,966 | 1.09% |
| Unclassified: |  | 777,756 | 194,902,752 | 37.18% |
| Total interspersed<br>repeats: |  |  | 403,732,765 | 77.02% |
| Small RNA: |  | 15,557 | 2,652,405 | 0.51% |
| Satellites: |  | 2,895 | 658,174 | 0.13% |
| Simple repeats: |  | - | - | 0.00% |
| Low complexity: |  | - | - | 0.00% |

\* most repeats fragmented by insertions or deletions have been counted as one element.

## Conclusions

This study presented a high quality genome assembly, annotation, and analysis of *M. nigrescens*. The genome was assembled using the long and linked-read sequencing data. Transcriptomic data from different developmental stages of the nematode was also generated. The *M. nigrescens* genome showed very high level of repeat content compared to other nematode genomes. These new resources and information will contribute to the better understanding of the genomic architecture and biology of mermithid nematodes and their adaptation to the broad range of insect the infect.

